# The Neuroprotective Effect of Short-chain Fatty Acids Against Hypoxia-reperfusion Injury

**DOI:** 10.1101/2023.12.23.573085

**Authors:** Anjit K. Harijan, Retnamony Kalaiarasan, Amit Kumar Ghosh, Ruchi P. Jain, Amal Kanti Bera

## Abstract

Gut microbe-derived short-chain fatty acids (SCFAs) are known to have a profound impact on various brain functions, including cognition, mood, and overall neurological health. However, their role, if any, in protecting against hypoxic injury and ischemic stroke has not been extensively studied. In this study, we investigated the effects of two major SCFAs abundant in the gut, propionate (P) and butyrate (B), on hypoxia-reperfusion injury using a neuronal cell line and a zebrafish model. Neuro 2a (N2a) cells treated with P and B exhibited reduced levels of mitochondrial and cytosolic reactive oxygen species (ROS), diminished loss of mitochondrial membrane potential, suppressed caspase activation, and lower rates of cell death when exposed to CoCl_2_-induced hypoxia, compared to the control group. Furthermore, adult zebrafish fed with SCFAs-supplemented feeds showed less susceptibility to hypoxic conditions compared to the control group, as indicated by multiple behavioral measures. Histological analysis of TTC-stained brain sections revealed lesser damage in the SCFAs-fed group. We also found that FABP7 (also known as BLBP), a neuroprotective fatty acid binding protein, was upregulated in the brains of the SCFAs-fed group. Additionally, when FABP7 was overexpressed in N2a cells, it protected the cells from hypoxia-reperfusion injury. Overall, our data clearly demonstrates a neuroprotective role of P and B against hypoxic brain injury and suggests the potential of dietary supplementation with SCFAs to mitigate stroke-induced brain damage.

**Highlights:**

- Short-chain fatty acid (SCFA) Propionate (P) and Butyrate (B) protect N2a cells from hypoxia-reperfusion.
- Zebrafish, when fed an SCFA-supplemented diet, are more resilient to hypoxia-reperfusion.
- SCFAs in the diet boost brain expression of FABP7 (fatty acid binding protein).
- FABP7 overexpression in N2a cells provides protection against hypoxia-reperfusion.
- SCFAs reduce reactive oxygen species (ROS) levels and increase FABP7, contributing to neuroprotection.

## 1. Introduction

Ischemic stroke, caused by the obstruction of blood flow to certain brain regions, affects millions of people worldwide. Despite current therapeutic options like thrombolytic drugs or thrombectomy, expected results are not always achieved. Moreover, the brain damage becomes irreversible if treatment is not provided within the critical time window. Since the last decade, the role of microbiota living in the human body has emerged as a critical determinant of brain functions, immune response, and overall health (Belkaid & Hand, 2014; Liu et al., 2022; Sasso et al., 2023). Notably, short-chain fatty acids (SCFAs), primarily acetate, propionate, and butyrate, are formed as metabolic byproducts during the fermentation of dietary fibers by gut microbes. These SCFAs have been shown to play a pivotal role in modulating brain function and behavior. Studies have demonstrated that post-stroke administration of sodium butyrate ameliorates brain inflammation in a carotid artery occlusion rat stroke model (Jaworska et al., 2017). Sodium butyrate also demonstrated a neuroprotective effect in a mouse model of traumatic brain injury (Li et al., 2016). There is ample evidence showing the protective role of SCFAs against a plethora of neurological disorders, such as Alzheimer’s disease, Parkinson’s disease, amyotrophic lateral sclerosis, depression, and other cognitive dysfunctions (Ameen et al., 2022; Duan et al., 2023; Mirzaei et al., 2021; Sun et al., 2023). The molecular mechanisms underlying neuroprotection caused by SCFAs are not yet fully understood. Numerous potential mechanisms have been proposed, and several studies have provided support for enhanced mitochondrial function, attenuation of oxidative stress, and promotion of the expression of various neuroprotective genes (Fock & Parnova, 2023; Mirzaei et al., 2021; Rose et al., 2018; van de Wouw et al., 2018). SCFAs, particularly butyrate, is known to inhibit histone deacetylases (HDACs), which could potentially contribute to its neuroprotective effects. Inhibition of HDACs leads to hyperacetylation of histones and subsequent remodelling of chromatin, making certain genes more accessible for transcriptional machinery and potentially increasing their expression (Candido, 1978; Davie, 2003). However, it is important to note that a recent report has challenged the popular view of HDAC inhibition by SCFAs. Instead, the report suggests that SCFAs activate the acetyltransferase p300, which is responsible for hyperacetylation (Thomas & Denu, 2021). SCFAs serve as ligands for a class of G protein-coupled receptors known as free fatty acid receptors (FFARs) (Offermanns, 2014). The activation of FFARs may mediate several functions of SCFAs (Falomir-Lockhart et al., 2019). Fatty acid binding proteins (FABPs) function as intracellular chaperones, playing a crucial role in regulating the transport, uptake, storage, and metabolism of lipophilic compounds, especially free fatty acids (Chmurzyńska, 2006; Storch & Corsico, 2023). FABPs may be associated with the effects mediated by SCFAs. Among the 12 members, FABP7 demonstrates high expression in neuronal tissue, particularly in oligodendrocytes and astrocytes (H. Kim et al., 2008; Matsumata et al., 2016). FABP7 has been implicated in playing a neuroprotective role, primarily by promoting neurogenesis, although some conflicting reports have raised doubts (Hamilton et al., 2023; Needham et al., 2022; Rui et al., 2019). In mouse astrocytes, FABP7 has been shown to offer protection against reactive oxygen species (ROS)-mediated toxicity via lipid droplet formation (Islam et al., 2019). Kato et al. investigated the role of FABP7 in ischemic brain injury (Kato et al., 2020). They subjected male mice to transient forebrain ischemia for 20 minutes and found that the level of injury did not vary significantly between wild-type (WT) and FABP7 knockout (KO) mice. However, compared to WT mice, KO mice exhibited a significant decrease in neurogenesis both in physiological and ischemic conditions. Interestingly, after 7-10 days of ischemia, the expression of FABP7 increased in neural stem/progenitor cells, coinciding with the peak of hippocampal neurogenesis (Kato et al., 2020). In contrast to this, Guo et al. reported that in mice a synthetic molecule MF6, which binds to FABP7 with high affinity and decreases the post-ischemic rise of FABP7, ameliorated brain injury caused by transient middle cerebral artery occlusion and reperfusion (Guo et al., 2021). Our present study investigates the potential of dietary supplementation of SCFAs to mitigate brain damage caused by hypoxia-reperfusion. In a zebrafish model, this supplementation yielded positive effects, and the underlying mechanisms appear to involve the upregulation of FABP7 and ROS scavenging.

## 2. Materials and methods

### 2.1. Reagents

Sodium propionate and sodium butyrate were obtained from Himedia Laboratories, India. Fluorescein diacetate (FDA), propidium iodide (PI), 2’,7’-dichlorodihydrofluorescein diacetate (DCFDA), MitoSOX Red, tetramethyl rhodamine ethyl ester perchlorate (TMRE), and MTT were purchased from Invitrogen. PCDNA3-FlipGFP (Casp3 cleavage seq) T2A mCherry obtained from Addgene (Catalogue ID 124428). Cobalt(II) chloride, and 2,3,5-Triphenyltetrazolium chloride (TTC) were purchased from Sigma Aldrich. FABP7 plasmid construct was a generous gift from Dr. Rosaline Godbout, University of Alberta, Canada.

### 2.2. Cell viability assay

Neuro 2a (N2a) cells were cultured in a cell culture incubator at 37°C with 5% CO_2_. Cells were seeded in a 35 mm dish with 3× 10^5^ cells and supplied with DMEM, supplemented with 10% FBS, and 1x penicillin and streptomycin. Cell viability was estimated by double staining with FDA and PI, which stain live and dead cells, respectively. For staining, the cells were incubated in serum-free DMEM, and FDA (50 nM) and PI (400 nM) were added. After 20 minutes of incubation at 37°C in the dark, the cells were washed with PBS and then imaged using an inverted fluorescent microscope (Olympus, IX83) with appropriate filter set and a sCMOS camera. The percent viability was calculated as [number of live cells / (number of dead cells + live cells)] x 100.

### 2.3. Measurement of reactive oxygen species (ROS) and mitochondrial potential **(ΔΨm)**

Intracellular ROS were measured with a redox probe DCFDA (10 µM), and mitochondrial ROS were measured with MitoSOX Red (5 µM), an indicator of superoxide. ΔΨm was estimated with TMRE (100 nM). Cells were incubated with the respective dyes in serum-free media at 37°C for 20 minutes. After washing with PBS, cells were imaged. Fluorescence intensity was measured offline after subtracting the background, with the help of software, ImageJ or CellSens (Olympus Corporation).

### 2.4. Caspase-3 activation

Caspase-3 activation was monitored using a genetically encoded reporter called ‘FlipGFP’ (PCDNA3-FlipGFP (Casp3 cleavage sequence) T2A mCherry (Q. Zhang et al., 2019). FlipGFP is a GFP-based fluorogenic protease reporter in which one of the beta strands flips in the GFP configuration. This beta strand is restored upon protease activation and cleavage, leading to the reconstitution of GFP and resulting fluorescence. Caspase-3 activation was monitored by measuring the relative intensity of green fluorescence to mCherry. An increase in green fluorescence indicates caspase-3 activation.

### 2.5. Hypoxia-reperfusion on N2a cells

Hypoxia was induced with CoCl_2_ (Gotoh et al., 2012; S. Zhang et al., 2011). Cells were incubated for 12 h at 37°C in serum-free DMEM, supplemented with 1 mM of CoCl_2_. For reperfusion, the media was replaced with CoCl_2_-free fresh media and incubated for another 6 h.

### 2.6. Animals and Housing

One hundred twenty adult wild-type zebrafish (short fin) were procured from a commercial supplier (Tarun Fish Farm, Chennai, India). These zebrafish were about 6 months old, and their gender remained undetermined. They were accommodated in 50-liter aquaria, maintained at an approximate temperature of 28°C, and the pH was regulated within the range of 7.0 to 8.0, under a 14 h:10 h light and dark photoperiod. Continuous aeration was provided to ensure adequate oxygen levels. They were fed with commercially available dry fish-food pellets (Optimum Micro Pellets) twice daily, with or without supplementation of SCFAs. The animals were allocated randomly into four groups, each consisting of 30 individuals: the control group (C), the group fed with sodium propionate (P), the group fed with sodium butyrate (B), and the group receiving a combination of both sodium propionate and sodium butyrate (P+B). The food pellets for each group were supplemented as follows: C: 1% NaCl; P: 1% sodium propionate; B: 1% sodium butyrate; P+B: a combination of 1% sodium propionate and 1% sodium butyrate. The feeding was continued for 2 months before conducting the hypoxia experiment.

### 2.7. Hypoxic treatment of zebrafish

The fish were exposed to hypoxia following the methods described earlier with slight modification (Braga et al., 2013; Das et al., 2019). In brief, a 1000 ml clear glass apparatus containing 800 ml of water was utilized (Fig. 2.a). An oximeter (Lutron DO-5509) was employed to continuously monitor the levels of dissolved oxygen in the water. As detailed in a prior study, the chamber had two openings: one was connected to a nitrogen cylinder, and the other remained in contact with the surrounding air (Braga et al., 2013). The induction of hypoxic conditions involved the infusion of pure nitrogen into the chamber until the dissolved oxygen levels in the water fell below 0.6 mg L^−1^, a range previously established for inducing short-term severe hypoxia in zebrafish (Das et al., 2019; Y. Lee et al., 2018). Subsequently, one fish at a time was promptly transferred from the adaptation chamber to the hypoxic chamber, and the tank was hermetically sealed, creating a closed, airtight system to induce hypoxia. Oxygen levels were meticulously maintained throughout the hypoxic period and remained constant during the trials. All hypoxia trials were recorded using a video camera (Panasonic HC-V270) and subsequently analyzed by two independent trained observers. The animals exhibited a consistent sequence of behaviors, which included four distinct stages of hypoxia. These phenotypes were categorized as follows: 1st stage, swimming at the water’s surface; 2nd stage, loss of posture; 3rd stage, maintenance of opercular beats with sporadic movements; and 4th stage, mortality. Once they reached the 3rd stage, they were immediately transferred to a recovery container in normoxic conditions, for observation and behavioral monitoring. After one hour in the recovery container and gaining normal body position for swimming, novel tank test was performed.

### 2.8. Novel tank test (NTT)

For assessing the anxiety-like behavior, NTT was performed following standard procedure (Cachat et al., 2010). The experiments were conducted in a nontoxic trapezoidal glass chamber, measuring 18 cm in height, 25 cm in length, and 5 cm in width. Individual fish was carefully selected from a holding tank and placed into the novel tank. Their behavior was promptly recorded over a period of 10 minutes using a Panasonic HC-V270 camera. Subsequently, the recorded data were imported into the ToxTrac application software (available at https://toxtrac.sourceforge.io/) and analyzed.

### 2.9. Histology

The extent of brain damage was assessed using TTC staining (Braga et al., 2013; Das et al., 2019). The animals were cryo-anesthetized by being placed in ice-cold water and euthanized by decapitation to remove the brain. The whole brains were immediately immersed in 1 ml of a 2% TTC solution in PBS (pH 7.4) at 37°C for 40 minutes. Subsequently, the TTC solution was removed, and the brains were washed with PBS (pH 7.4). They were briefly stored at -20°C before being sliced. Images were taken with an upright microscope and analyzed.

### 2.10. Western blot

Whole brain was homogenized in RIPA lysis buffer (10 mM Tris-HCl, pH 7.5 containing 150 mM NaCl, 1% Nonidet P40, 0.1% SDS, 1% Triton X-100, PMSF 0.1 mg/mL) supplemented with proteinase and phosphatase inhibitors cocktail (1:100; Sigma Aldrich). Lysates were centrifuged at 13,000× *g* for 20 min at 4°C. The supernatant was collected and protein concentration was estimated with a Bio-Rad DC^TM^ protein assay kit. Total extracts were resolved on SDS-PAGE followed by transfer onto a PVDF membrane. The membranes were blocked for 1 h in Tris-buffered saline (TBS) solution containing 0.1% Tween (TBS-Tween) and 5% BSA, followed by overnight incubation with specific primary antibodies at 4°C with gentle agitation. Primary antibody of anti-FABP7 raised in rabbit (Ab32423; Abcam), and anti-β-actin raised in mouse (A5316; Sigma-Aldrich) were used in 1:1000 and 1:5000 dilutions respectively. After washing in TBS-Tween the blot was probed with HRP-conjugated secondary antibody. Immunoblot signals were visualized using an ECL chemiluminescence kit, captured with ChemiDoc^TM^ XRS and analysed using Quantity One software (Bio-Rad) or ImageJ.

### 2.11. MTT assay

The protective role of FABP7 against hypoxia-reperfusion injury was assessed by measuring cell viability with the MTT assay. 50 µl of a 5 mg/mL MTT solution in sterile PBS was added to each dish (1.0x 10^4^ cells/dish) and incubated for 3 h in a cell culture incubator. The medium was then removed, and MTT formazan crystals formed by the cells were dissolved using 150 µl of dimethyl sulfoxide (DMSO) for an additional 10 minutes in the dark. The OD in each well was determined using a microplate spectrophotometer (Bio-Rad Laboratories Inc.) at 490 nm. Cell viability was expressed as the percentage of MTT reduction, with the absorbance of the control cells assigned a value of 100%.

### 2.12. RT-qPCR

At the end of the behavioral study, zebrafish were euthanized by rapid chilling with submersion in an ice-water bath for 10-20 min. The whole brain was excised at 4°C. Three brains pool together and kept in a single tube. Total RNA was extracted with TRIzol reagent (Life Technologies) following the manufacturer’s protocols, and quantified using a Nanodrop 2000 (Thermo Fisher Scientific). The cDNA was transcribed from 300 ng total RNA using the reverse transcription system (Promega, USA) in a final volume of 50 μl. SYBR-Green-based real-time quantitative PCR was performed in an ABI QuantStudio 7 Flex machine (Applied Biosystems, USA). Fold-change was calculated by the delta-delta Ct method (Livak & Schmittgen, 2001). The final results were expressed as fold changes between the sample and the controls, corrected with housekeeping gene GAPDH. Primer sequences that were used for the experiments are listed in Table 1.

**Table 1.**
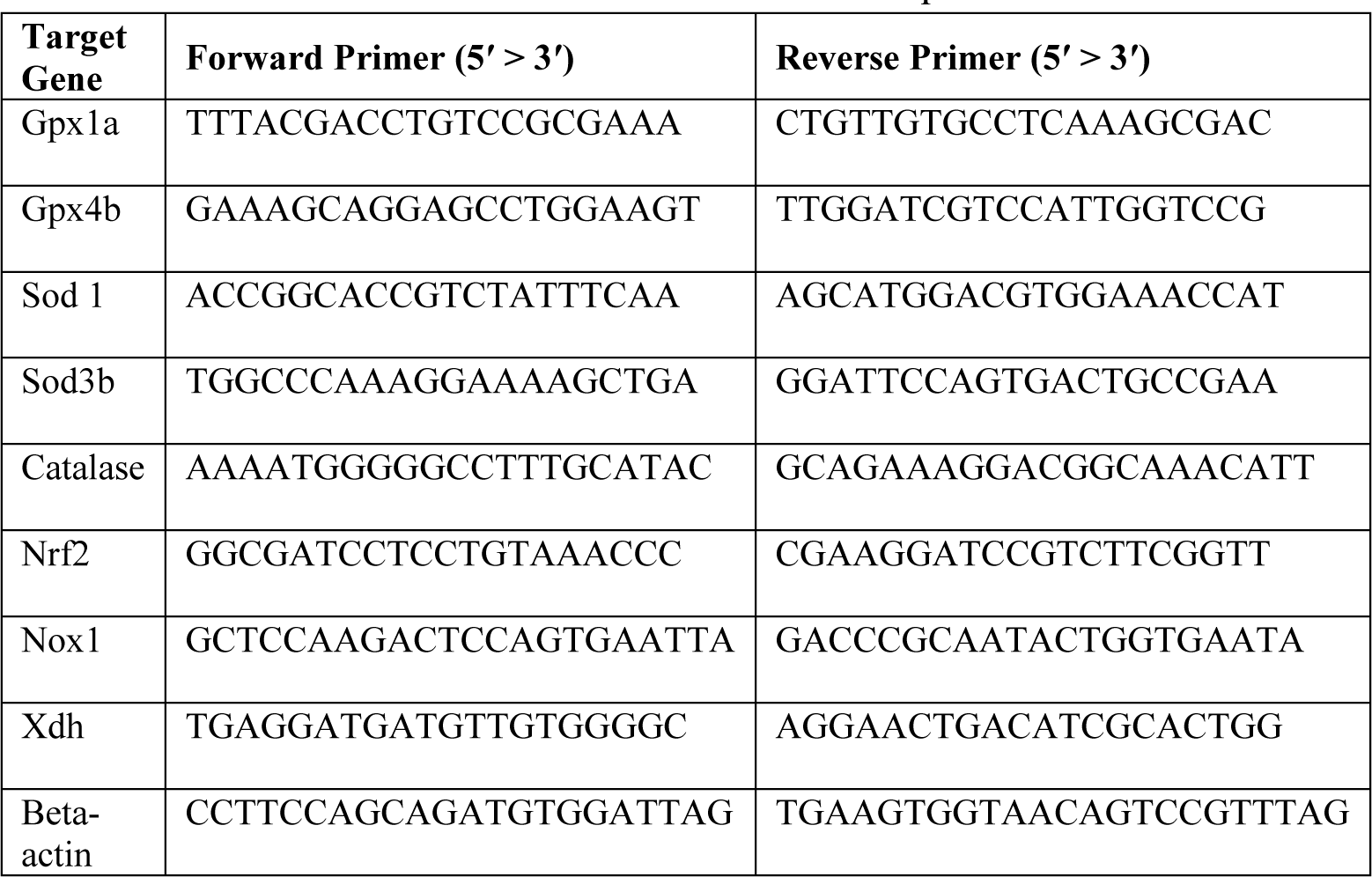
Primers used for the RT-qPCR.

### 2.13. Ethics Statement

All animal experiments were performed in accordance with the guidelines issued by the Institutional Animal Care and Use Committees.

### 2.14. Statistical analysis

Data are expressed as the mean ± standard error of the mean. Student’s t-test was used to compare between two groups. For multiple-group comparisons, one-way ANOVA was employed and Tukey’s post hoc test was used. Analyses were performed using the GraphPad Prism software, version 6.0.

## 3. Results

### 3.1. SCFAs prevent cell death from hypoxia-reperfusion injury

N2a cells exposed to CoCl_2_-induced hypoxia for 12 h and subsequent re-oxygenation with fresh media for 6 h showed significant cell death, as assessed with FDA/PI staining (Fig. 1a and S1a.). Interestingly, cells pretreated with P, B or P+B for 24 h improved the cell viability significantly. As shown in Fig.1a. cell viability reduced to 68 ± 8% upon hypoxia-reperfusion. P, B, and P+B treated cells showed viability of 95 ± 5%, 106 ± 6% and 109 ± 4% respectively, after hypoxia-reperfusion. These findings show that SCFA pre-incubation helps in the survival of cells under hypoxia-reperfusion in vitro.

**Fig. 1.**
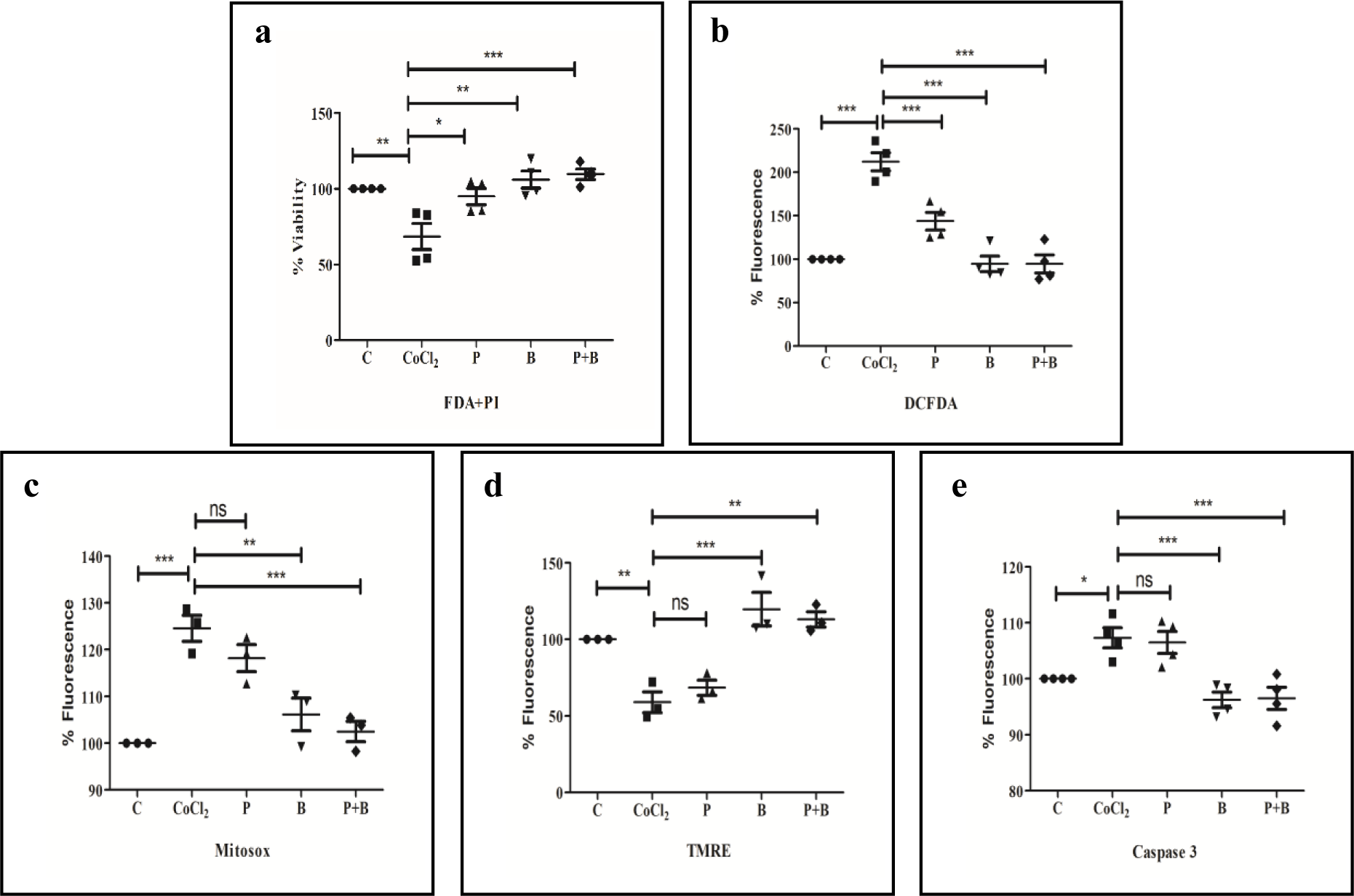
SCFAs mitigate oxidative stress, stabilize mitochondrial membrane potential, inhibit caspase-3 activation, and prevent cell death induced by hypoxia-reperfusion. **a**. FDA-PI double stain used to measure cell viability in Neuro-2a cells. Both propionate (P) and butyrate (B), individually and in combination, significantly reduce cell death. **b.** Intracellular ROS, measured with DCFDA significantly decreased in P, B and P+B treated cells. **c**. Mitochondrial superoxide levels quantified with Mitosox Red, in B and P+B treated cells are comparable to control cells. P alone failed to mitigate the rise in mitochondrial superoxide caused by CoCl_2_ treatment. **d**. CoCl_2_ treated cells show loss of ΔΨm, which is not present in B and P+B treated cells stained with TMRE **e**. Caspase-3 activation, monitored with FlipGFP. CoCl_2_ treatment increased caspase-3 activation, and treatment with P had no effect. The B and P+B treatment prevented caspase-3 activation caused by CoCl_2_. All the statistics obtained are from 3-4 independent experiments and a total of 200-400 cells in each group.

#### 3.1.1. Intracellular ROS

Cell death caused by hypoxia-reperfusion is typically preceded by a surge in ROS. We measured the ROS level using the ROS-sensitive dye DCFDA. As depicted in Fig. 1b and S1b, the ROS levels increased, as indicated by the elevated fluorescence, following hypoxia-reperfusion, and this increase was significantly attenuated in the SCFA-treated groups. In control cells, there was approximately a two-fold increase in fluorescence levels after hypoxia-reperfusion (Fig. 1b). The P treated group exhibited a 1.44-fold increase, whereas there was no significant rise in ROS levels in the B and P+B treated groups following hypoxia-reperfusion.

##### Mitochondrial ROS

Mitochondrial ROS (mito-ROS) was measured with MitoSOX Red. Fig.1c and S1c show mito-ROS in different experimental conditions. As expected, the CoCl_2_ treated group showed a higher level of mito-ROS (125 ± 3%) compared to the untreated control. However, P treatment couldn’t prevent the rise of mito-ROS (118 ± 3%). In contrast, B alone or in combination with P prevented the surge of mito-ROS significantly. The relative fluorescence intensities for B and P+B treated groups were 106 ± 4% and 103± 2% respectively.

#### 3.1.2. Mitochondrial potential (ΔΨm)

The loss of ΔΨm is a hallmark of hypoxia-reperfusion injury that results in cell death (Ashok et al., 2023; Weinberg et al., 2000). ΔΨm, measured with TMRE, revealed that N2a cells exposed to hypoxia-reperfusion had a lower ΔΨm compared to the untreated control (Fig. 1d and S1d). CoCl_2_ treatment significantly reduced TMRE fluorescence to 59 ± 3%, indicating mitochondrial depolarization. The B alone or in combination with P completely prevented the loss of ΔΨm. Although hypoxia-reperfusion caused a lesser loss of ΔΨm (69 ±5%) in P-treated cells, the result was not statistically significant (Fig. 1d).

#### 3.1.3. Caspase-3 activation

Activation of Caspase-3 has often been associated with various cell death processes (Eskandari & Eaves, 2022; Espinosa-Oliva et al., 2019). We assessed caspase-3 activation using a genetically encoded sensor called FlipGFP, as described in the methods section. Fig. 1e and S1e depict a significant increase of Caspase-3 activation following hypoxia-reperfusion. Although P couldn’t prevent caspase activation, B and P+B treated cells show no significant rise of Caspase-3 activities following CoCl_2_ treatment.

### 3.2. Effect of SCFAs on hypoxia-induced behavior of zebrafish

#### 3.2.1. SCFAs fed zebrafish recovered faster after hypoxia

To assess the protective effect of SCFAs against hypoxia, adult zebrafish were transferred to a hypoxic chamber, and the time it took them to reach stage 3 was recorded (Fig. 2a; see the methods). Both the control and SCFAs-fed groups took 6 to 7 minutes to reach stage 3, and there was no statistically significant difference among the different groups (data not shown). Further, we recorded the recovery time, that is, the time taken to regain normal body position for swimming after hypoxic shock. As shown in Fig. 2b zebrafish fed with an SCFAs-containing diet recovered faster compared to the control group. The control group took 17 ± 1.3 minutes, while the recovery times were 13 ± 0.6, 11.5 ± 0.7, and 12 ± 0.5 minutes for the P, B, and P+B fed groups, respectively.

**Fig. 2.**
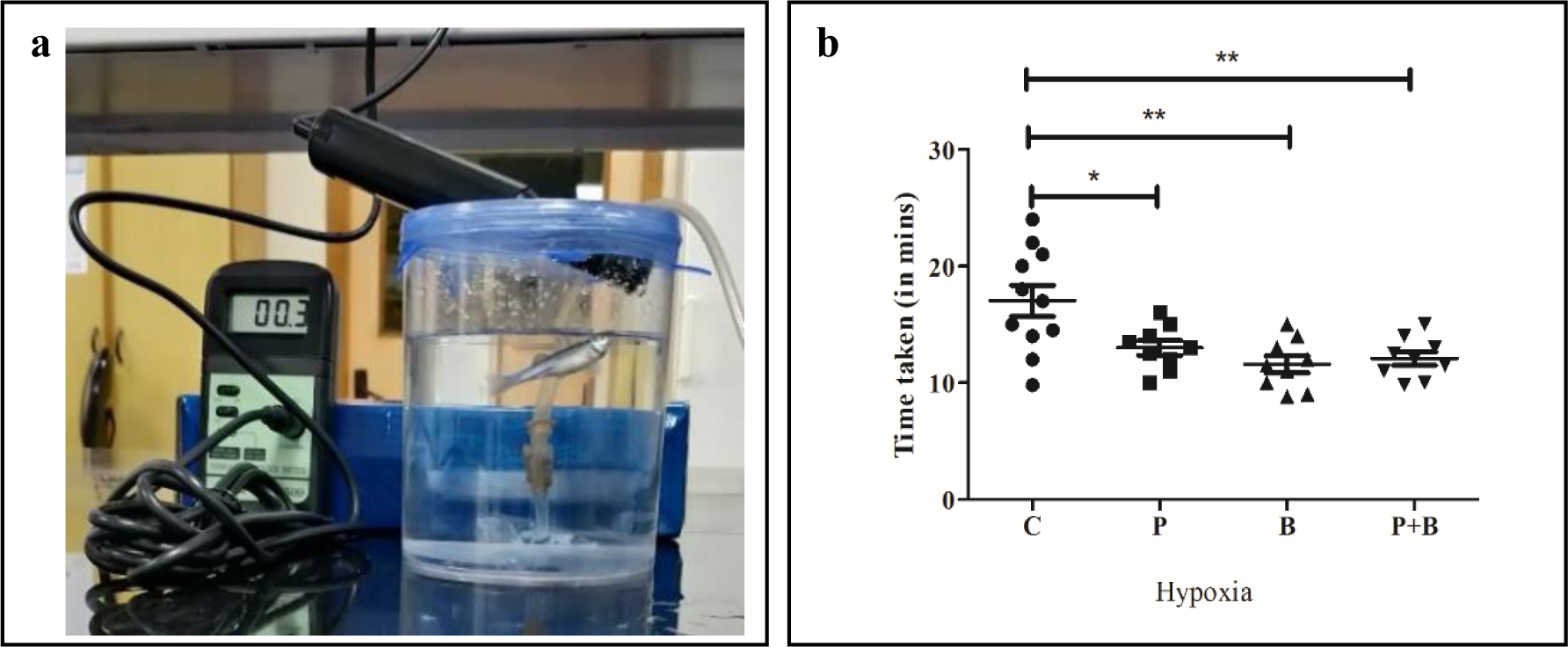
**a.** Image of dissolved oxygen meter used to monitor the dissolved oxygen level in hypoxia chamber after N_2_ gas purging. **b**. Recovery time after a hypoxic challenge is significantly reduced in fish fed with SCFAs.

#### 3.2.2. Locomotor activity and anxiety-like behavior

##### 3.2.2.1. Total distance travelled

Locomotor activity and anxiety-like behavior of the fish were assessed using the novel tank test (NTT), as described in the methods section. Swimming trajectories of adult zebrafish in normoxia and post-hypoxia are depicted in Fig. 3a and 3b. As evident in Fig. 3a, fish, if not challenged with hypoxia, exhibited random swimming patterns throughout the entire novel tank, regardless of whether they were fed a control diet or a diet supplemented with SCFAs. Over a 10-minute period, they covered a mean distance ranging from 70 meters to 125 meters, and there were no statistically significant differences between the control group and the SCFAs-fed groups (Fig. 3c). However, when the fish were exposed to hypoxia, their swimming pattern underwent a drastic change. They spent most of their time swimming near the water’s surface, and there was a significant reduction in the total distance traveled (Fig. 3b, 3c). Interestingly, following hypoxic treatment, the SCFAs-fed groups showed a trend of covering greater distances within the novel tank compared to the control group, which is an indication of resilience, although these differences did not reach statistical significance except the P+B group (68 ± 11 meters, p<0.05) (Fig. 3b, 3c).

**Fig. 3.**
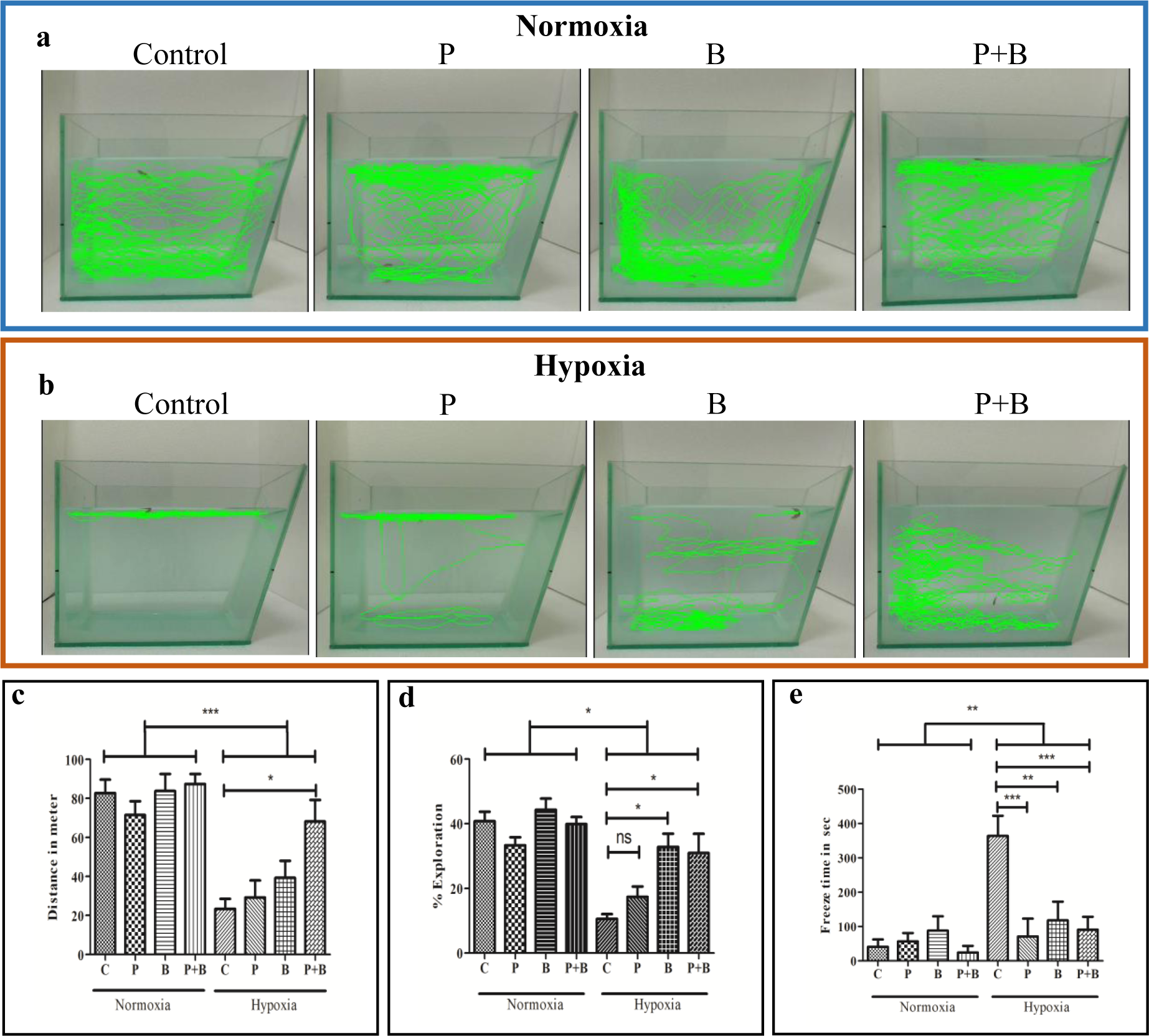
Swimming trajectory of zebrafish in a novel tank. Zebrafish fed with Propionate (P), Butyrate (B), and P+B containing diets exhibit different swimming patterns following a hypoxic challenge, compared to control. **a**. Representative swimming trajectory of fish fed with control and SCFAs containing diet, under normoxia. **b**. Swimming trajectory following hypoxic challenge. **c.** Hypoxia diminishes locomotor activities. Locomotion in SCFAs-fed groups is less affected compared to the control group. After hypoxia, the P+B fed group covered the maximum distance, indicating a lesser impact of hypoxia. **d.** Percentage of the whole arena explored. Fish exposed to hypoxia explored a smaller area of the arena than the normoxic control. SCFAs-fed groups showed improved exploration. **e.** Hypoxia significantly increased freezing time. SCFAs supplementation reduced freezing time following a hypoxic insult. Values are expressed as mean ± SEM. Each group contains 9-12 fishes. Statistical significance is denoted as *p<0.05, **p<0.01, ***p<0.001, ****p<0.0001. ‘ns’ indicates not significant.

##### 3.2.2.2. Exploratory behavior

We measured the percentage of the total tank arena explored by the fish. The exploration of a larger area reflects the zebrafish’s curiosity about its environment under normal conditions. Typically, stressed or anxious fish do not explore their arena as much. As shown in Fig. 3a, 3d under normoxia, approximately 41±3% of the arena was explored by both the control and SCFAs-fed fish. After hypoxia, the tank arena exploration significantly decreased for both the control and SCFAs-fed groups (Fig. 3b, 3d). However, the SCFAs-fed group explored a larger area of the tank post-hypoxia compared to the control diet-fed group (control, 11 ± 2%; P, 17 ± 3%; B, 33 ± 4%; and P+B, 31 ± 6%) (Fig. 3d), implying that SCFAs-fed groups are more resilient to hypoxia-induced stress.

##### 3.2.2.3. Freezing time

We estimated freezing time in NTT to assess the level of fear/stress in zebrafish post-hypoxia based on a 10-minute recording. As shown in Fig. 3e, freezing time significantly increased in hypoxia-exposed groups compared to the control (Fig. 3e). In accordance with other behavioral parameters, SCFAs-fed groups were immobilized for a shorter duration compared to the control group (control, 364 ± 59 sec; P, 71 ± 52 sec; B, 118 ± 54 sec; P+B, 91 ± 38 sec), further emphasizing the protective role of SCFAs.

### 3.3. SCFAs protect the brain from hypoxic damage

The extent of brain injury was assessed with TTC staining. TTC-stained healthy brain sections and hypoxic-damaged brain sections are shown in Fig. 4. As evident in the Figure, healthy brains exhibited deep red staining bilaterally, while post-hypoxia brains displayed several infarcts as pale or unstained regions spreading from the ventro-dorsal region to the deep structure of the midbrain (Fig. 4a and b). The infarct areas calculated with ImageJ revealed significantly smaller infarct size in the SCFAs-fed fish brain compared to the normal diet-fed fish. Infarct size, measured as a percentage of the total brain area, was normalized against the respective normoxic control and presented in Fig. 4c.

**Fig. 4.**
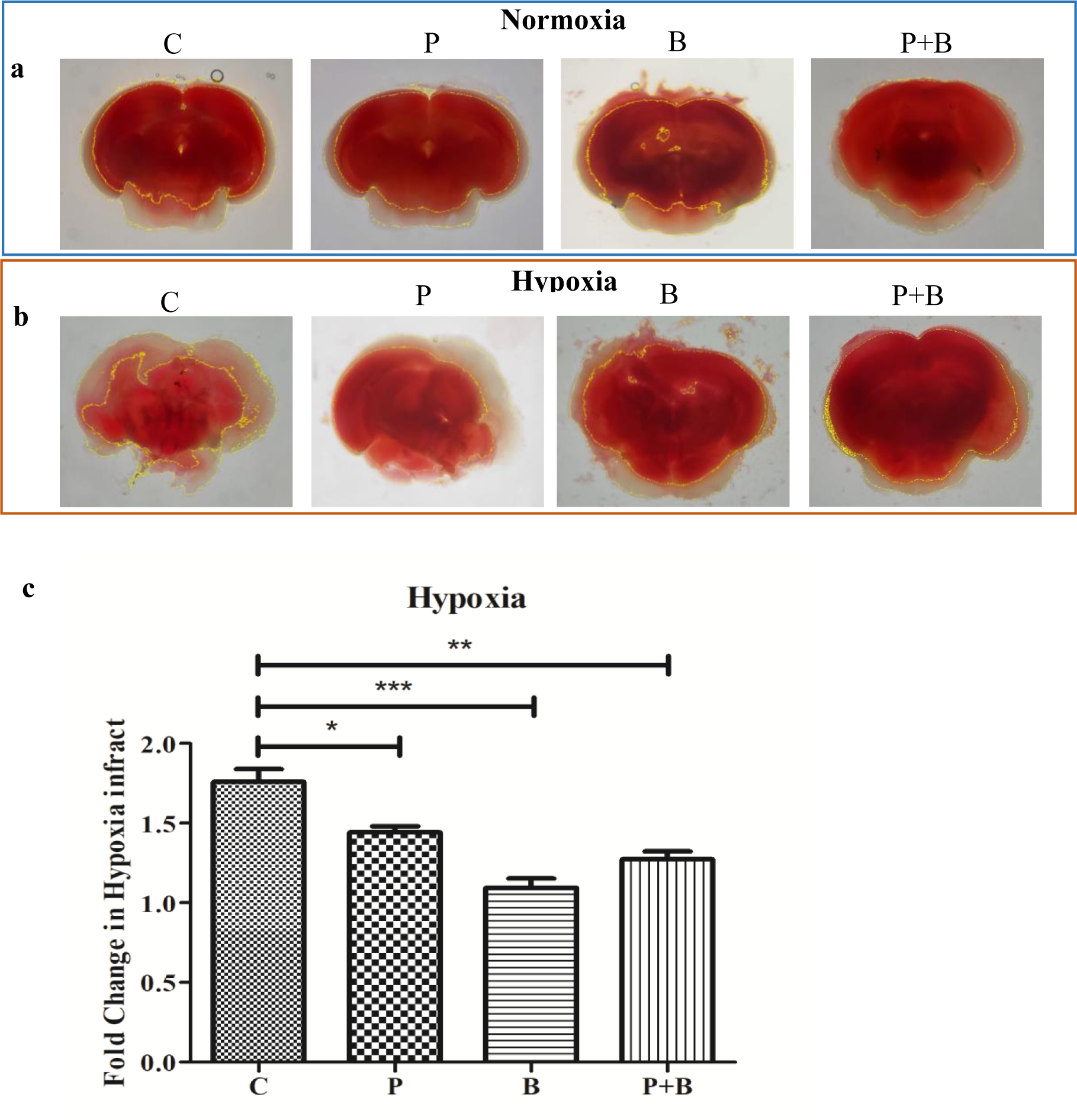
Hypoxia-induced brain damage, evaluated with TTC staining. **a**. Images of control and SCFAs-fed zebrafish brain slices in normoxic conditions. TTC staining shows negligible infarction. **b**. Infarction caused by hypoxia in control and SCFAs-fed fish brain sections. **c**. Statistics showing reduced infarction in the SCFAs-fed group. Total infarct area in hypoxic brain slices is normalized against corresponding normoxic brain sections. Each group contains data from 5-8 fishes. *p< 0.05, **p<0.01, ***p<0.001. P: propionate, B: butyrate.

### 3.4. Expression of Pro- and antioxidant genes

The expression levels of various pro- and antioxidant genes in the fish brain were compared by measuring mRNA levels using real-time PCR (Fig. 5). The expression levels in the groups fed with SCFAs were compared to those in the group fed with the control diet. ROS scavenging enzymes, superoxide dismutase (SOD) 1 and 3b, were upregulated in all SCFAs-fed groups (P, B, and P+B). The expression of two other antioxidant enzymes, catalase and glutathione peroxidase (GPX) 4b, also significantly increased in all SCFAs-fed groups. Also, there was a significant upregulation of antioxidant gene GPX1a in the P+B fed group. Although not statistically significant, both the P and B fed groups showed a trend of increased GPX1a expression. We studied the expression of a transcription factor, NRF2, which is considered as a master regulator of several antioxidant genes. It was found to be significantly increased in the B and P+B fed groups. Furthermore, we measured the levels of mRNAs of ROS-producing enzymes NADPH oxidase (Nox) and xanthine oxidase (XDH). Nox1 and XDH decreased significantly in all SCFAs-fed groups (Fig. 5).

**Fig. 5.**
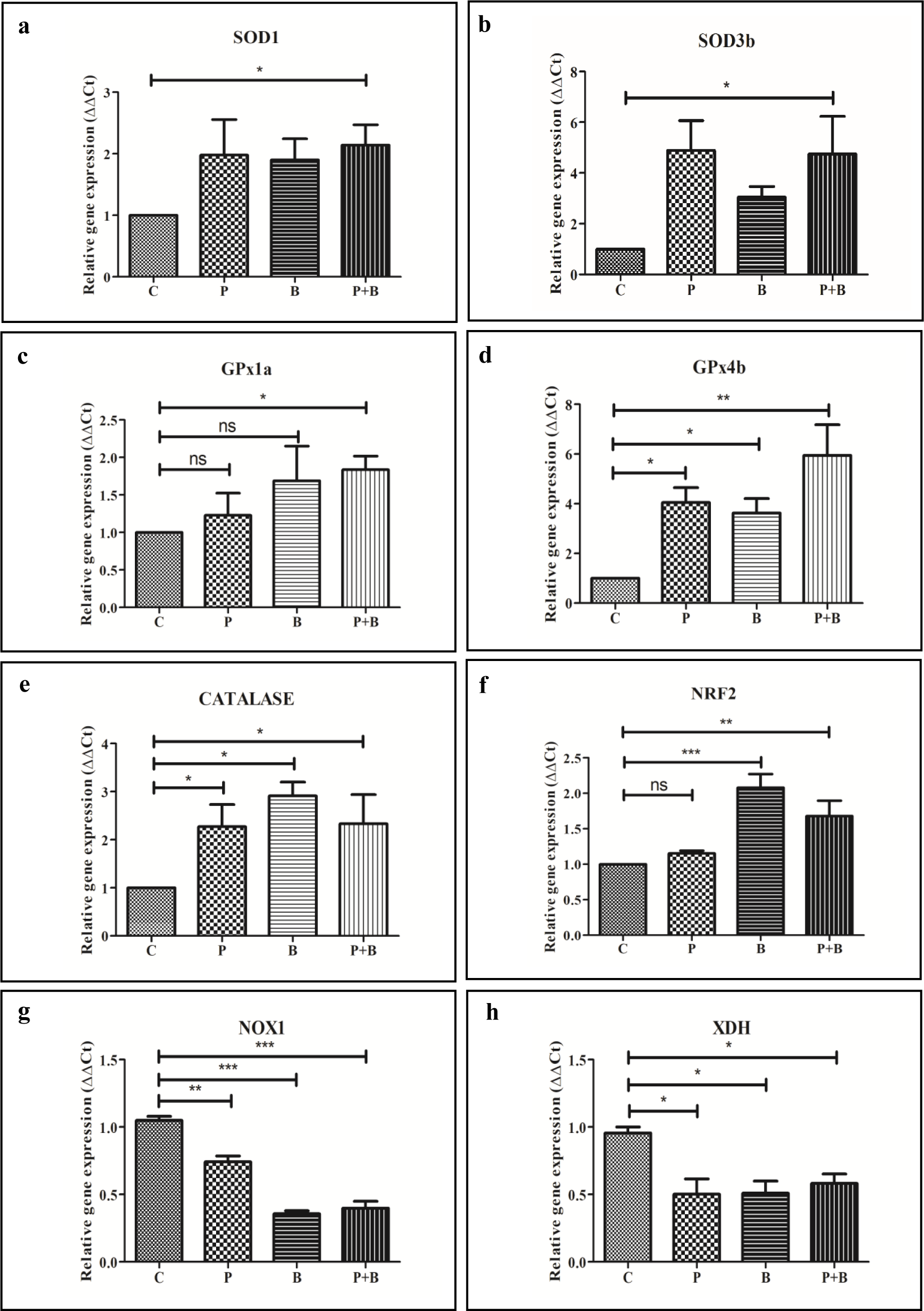
Real-time quantitative PCR analysis of zebrafish brain. **a-b**. SOD1 and SOD3b mRNA expression increased significantly in SCFAs-fed fish. **c**. GPX1a expression increased significantly in the P+B-fed group. The P and B-fed group showed a trend of increased expression, though it was not statistically significant. **d**. GPX4b increased significantly in all SCFA-fed groups. **e**. Catalase expression significantly increased following SCFAs feeding. **f**. Both B and P+B increased NRF2 expression. **g-h**. NOX1 and XDH1 expression decreased significantly upon SCFAs feeding. Values are expressed as mean ± SEM of three independent experiments. *p < 0.05, **p < 0.01, ***p < 0.001, ns: not significant.

### 3.5. Expression of Fatty acid binding protein 7 (FABP7)

FABP7 has been implicated in neurogenesis and neuroprotection (Foerster et al., 2020; Islam et al., 2019). We investigated its potential involvement in the protective role of SCFAs against hypoxia-reperfusion injury. Western blot analysis of brain tissues, as shown in Fig. 6a-c, reveals that SCFAs-fed groups exhibit higher levels of FABP7 protein. In comparison to the control group fed a normal diet, FABP7 expression was 1.4 ± 0.1, 2.2 ± 0.2, and 2.5 ± 0.3-fold higher in the P, B, and P+B-fed groups, respectively (Fig. 6b). Furthermore, challenging with hypoxia did not result in any further increase in FABP7 levels in the SCFAs-fed groups (Fig. 6c). Interestingly, in contrast to the SCFAs-fed groups, fish fed with the control diet showed an increased expression of FABP7 following hypoxia. We conducted a similar study in the N2a cell line. Consistent with the zebrafish results, cells treated with B and P+B showed increased expression of FABP7, with folds of 1.24 and 1.44, respectively (Fig. 6d and 6e). While we occasionally observed increased FABP7 levels following propionate treatment, the cumulative data did not reach statistical significance. Chemically induced hypoxia with CoCl_2_ did not significantly increase FABP7 expression in all groups except for the P+B-treated group. Cells treated with P+B exhibited a 1.9-fold increase in FABP7 expression after hypoxia-reperfusion (Fig. 6d and 6f).

**Fig. 6.**
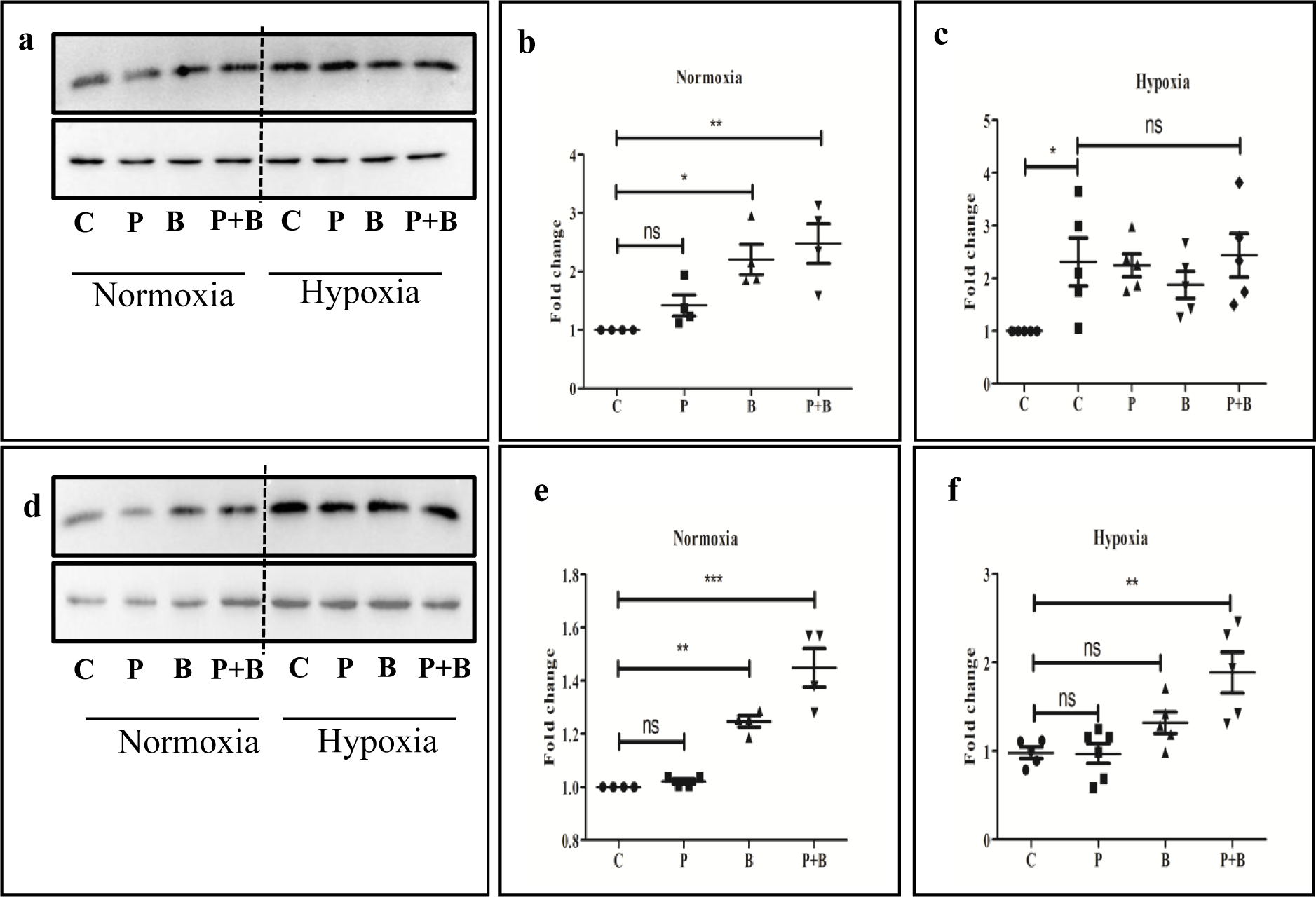
**a**. Western blot showing FABP7 expression in the zebrafish brain. Homogenates of 3 pooled brain samples were loaded on each lane. **b.** Data summary shows increased expression of FABP7 in the B and P+B fed groups. **c**. Hypoxia significantly increased FABP7 levels in the brains of fish under the control diet but not in the SCFAs-fed groups. Data presented in Fig. b and c are extracted from multiple blots like the one presented in Fig. **a**. **d**. Expression of FABP7 in Neuro-2a cells. Statistics derived from multiple blots are shown in Fig. e and f. **e**. B and P+B increased FABP7 expression in Neuro-2a cells. **f**. Hypoxia increased FABP7 expression in P+B treated cells. Hypoxia was induced with 1 mM CoCl_2_. Values are expressed as mean ± SEM. *p < 0.05, **p < 0.01, ***p < 0.001, ns - not significant.

### 3.6. Overexpression of FABP7 improves cell viability in hypoxia-reperfusion

N2a cells were transfected with a plasmid encoding human FABP7, while mock-transfected control cells received an eGFP plasmid. After 12 h of CoCl_2_-induced hypoxia and 6 h of reperfusion, cell viability was assessed using the MTT assay. As shown in Figure 7, FABP7 overexpression significantly improved cell viability.

**Fig. 7.**
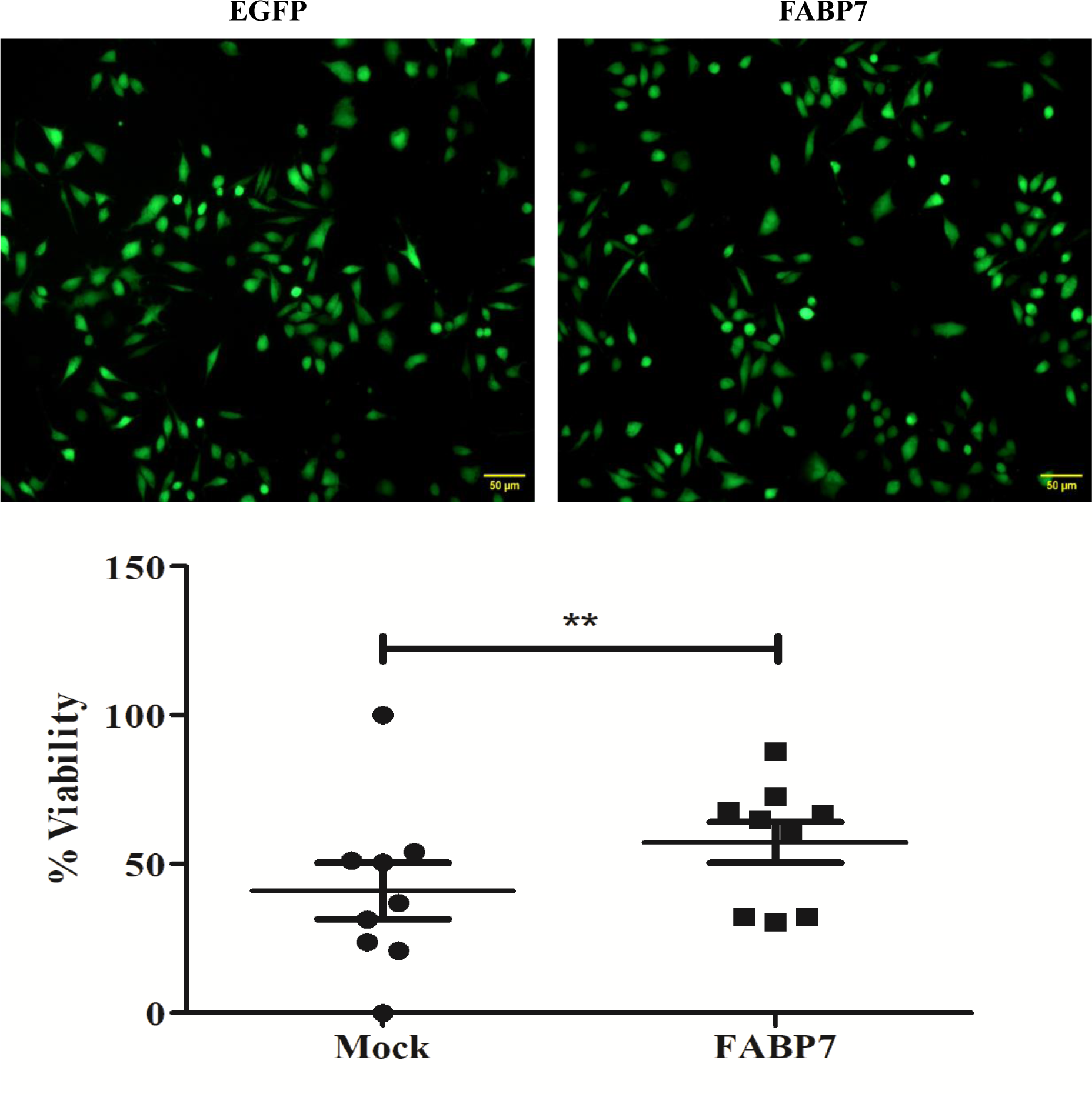
Overexpression of FABP7 protects N2A cells from hypoxia-reperfusion injury. Images depict the expression of EGFP (control) and EGFP-tagged FABP7. After inducing hypoxia with CoCl_2_ followed by reperfusion with fresh media, cell viability was estimated using an MTT assay. FABP7-overexpressed cells exhibited significantly higher viability. Values are expressed as mean ± SEM. **p < 0.01.

## 4. Discussion

In this study, we have demonstrated that SCFAs, namely propionate (P) and butyrate (B), either individually or in combination, are capable of reducing cell death induced by hypoxia-reperfusion. Zebrafish that were fed with a diet supplemented with P and B for two months exhibited significant resilience, faster post-stroke recovery, reduced brain infarction, and improved behavioral outcomes. The underlying mechanisms appear to involve the attenuation of ROS levels due to alterations in the expression of genes associated with ROS homeostasis. When compared to P, B was found to be more effective in certain experimental setups, and combining P with B did not provide any additional benefits over using B alone. Our data on remodelling of ROS signalling pathways by SCFAs align with previous studies conducted on different cell lines by other research groups (González-Bosch et al., 2021; Huang et al., 2017; S. Y. Kim et al., 2020; Robles-Vera et al., 2020). In fact, ROS is believed to play a central role in various neurological disorders such as stroke, Alzheimer’s disease, and Parkinson’s disease, where dysregulation of the gut-brain axis has been identified as a contributing factor (Cryan et al., 2019). In a recent study, Lee et al. observed reduced levels of SCFAs in the feces of aged mice compared to younger ones, and recolonizing older mice with SCFA-producing bacteria led to better outcomes in post-stroke recovery (J. Lee et al., 2020). Alongside the reduction in cytoplasmic and mitochondrial ROS, SCFA treatment resulted in less dissipation of ΔΨm and decreased caspase activation in cells exposed to hypoxia-reperfusion. This suggests that SCFAs assist in maintaining cellular redox balance, preventing cells from undergoing apoptosis. Furthermore, we report a novel finding: an increase in the expression of FABP7 due to SCFAs and its potential contribution to preventing hypoxia-reperfusion injury. Fish fed with SCFAs exhibited higher levels of FABP7 protein in their brains. Interestingly, FABP7 expression in the brains of control fish increased to the same level as the SCFAs-fed group after exposure to hypoxia. Hypoxia did not further increase FABP7 levels in the SCFAs-fed group, which were already elevated. Presumably, in control fish, the increase in FABP7 expression after hypoxia aims at brain repair by promoting neurogenesis, while overcoming a hypoxic challenge requires a higher baseline level of FABP7. When FABP7 was overexpressed in N2a cells, it improved cell viability during hypoxia-reperfusion, validating its positive contribution to stroke recovery.

In addition to stroke, many neurodegenerative diseases, such as Alzheimer’s and Parkinson’s disease, involve significant loss of neurons. Stimulating neurogenesis through increased FABP7 levels with dietary supplementation of SCFAs may be a promising approach for enhancing therapeutic outcomes.

In conclusion, our study demonstrates the protective effects of SCFAs against hypoxia-reperfusion injury. The underlying mechanisms include the reduction of ROS levels and the upregulation of a protein, FABP7, which is associated with neurogenesis.

## Acknowledgements

The work was supported by the Indian Council of Medical Research. AKH received a fellowship from the University Grants Commission. We acknowledge Dr. Gomathy and Athira for their critical comments on the manuscript.

## Declaration of Generative AI and AI-assisted technologies in the writing process

During the preparation of this work the authors used ChatGPT in order to improve readability and language. After using this tool, the authors reviewed and edited the content as needed and take full responsibility for the content of the publication.

## References

Ameen, A. O., Freude, K., & Aldana, B. I. (2022). Fats, Friends or Foes: Investigating the Role of Short- and Medium-Chain Fatty Acids in Alzheimer’s Disease. Biomedicines, 10(11), 2778. 10.3390/biomedicines10112778

Ashok, D., Papanicolaou, K., Sidor, A., Wang, M., Solhjoo, S., Liu, T., & O’Rourke, B. (2023). Mitochondrial membrane potential instability on reperfusion after ischemia does not depend on mitochondrial Ca2+ uptake. Journal of Biological Chemistry, 299(6), 104708. 10.1016/j.jbc.2023.104708

Belkaid, Y., & Hand, T. W. (2014). Role of the Microbiota in Immunity and Inflammation. Cell, 157(1), 121–141. 10.1016/j.cell.2014.03.011

Braga, M. M., Rico, E. P., Córdova, S. D., Pinto, C. B., Blaser, R. E., Dias, R. D., Rosemberg, D. B., Oliveira, D. L., & Souza, D. O. (2013). Evaluation of spontaneous recovery of behavioral and brain injury profiles in zebrafish after hypoxia. Behavioural Brain Research, 253, 145–151. 10.1016/j.bbr.2013.07.019

Cachat, J., Stewart, A., Grossman, L., Gaikwad, S., Kadri, F., Chung, K. M., Wu, N., Wong, K., Roy, S., Suciu, C., Goodspeed, J., Elegante, M., Bartels, B., Elkhayat, S., Tien, D., Tan, J., Denmark, A., Gilder, T., Kyzar, E., … Kalueff, A. V. (2010). Measuring behavioral and endocrine responses to novelty stress in adult zebrafish. Nature Protocols, 5(11), 1786–1799. 10.1038/nprot.2010.140

Candido, E. (1978). Sodium butyrate inhibits histone deacetylation in cultured cells. Cell, 14(1), 105–113. 10.1016/0092-8674(78)90305-7

Chmurzyńska, A. (2006). The multigene family of fatty acid-binding proteins (FABPs): Function, structure and polymorphism. Journal of Applied Genetics, 47(1), 39–48. 10.1007/BF03194597

Cryan, J. F., O’Riordan, K. J., Cowan, C. S. M., Sandhu, K. V., Bastiaanssen, T. F. S., Boehme, M., Codagnone, M. G., Cussotto, S., Fulling, C., Golubeva, A. V., Guzzetta, K. E., Jaggar, M., Long-Smith, C. M., Lyte, J. M., Martin, J. A., Molinero-Perez, A., Moloney, G., Morelli, E., Morillas, E., … Dinan, T. G. (2019). The Microbiota-Gut-Brain Axis. Physiological Reviews, 99(4), 1877–2013. 10.1152/physrev.00018.2018

Das, T., Soren, K., Yerasi, M., Kumar, A., & Chakravarty, S. (2019). Revealing sex-specific molecular changes in hypoxia-ischemia induced neural damage and subsequent recovery using zebrafish model. Neuroscience Letters, 712, 134492. 10.1016/j.neulet.2019.134492

Davie, J. R. (2003). Inhibition of Histone Deacetylase Activity by Butyrate. The Journal of Nutrition, 133(7), 2485S–2493S. 10.1093/jn/133.7.2485S

Duan, W.-X., Wang, F., Liu, J.-Y., & Liu, C.-F. (2023). Relationship Between Short-chain Fatty Acids and Parkinson’s Disease: A Review from Pathology to Clinic. Neuroscience Bulletin. 10.1007/s12264-023-01123-9

Eskandari, E., & Eaves, C. J. (2022). Paradoxical roles of caspase-3 in regulating cell survival, proliferation, and tumorigenesis. Journal of Cell Biology, 221(6). 10.1083/jcb.202201159

Espinosa-Oliva, A. M., García-Revilla, J., Alonso-Bellido, I. M., & Burguillos, M. A. (2019). Brainiac Caspases: Beyond the Wall of Apoptosis. Frontiers in Cellular Neuroscience, 13. 10.3389/fncel.2019.00500

Falomir-Lockhart, L. J., Cavazzutti, G. F., Giménez, E., & Toscani, A. M. (2019). Fatty Acid Signaling Mechanisms in Neural Cells: Fatty Acid Receptors. Frontiers in Cellular Neuroscience, 13. 10.3389/fncel.2019.00162

Fock, E., & Parnova, R. (2023). Mechanisms of Blood–Brain Barrier Protection by Microbiota-Derived Short-Chain Fatty Acids. Cells, 12(4), 657. 10.3390/cells12040657

Foerster, S., Guzman de la Fuente, A., Kagawa, Y., Bartels, T., Owada, Y., & Franklin, R. J. M. (2020). The fatty acid binding protein FABP7 is required for optimal oligodendrocyte differentiation during myelination but not during remyelination. Glia, 68(7), 1410–1420. 10.1002/glia.23789

González-Bosch, C., Boorman, E., Zunszain, P. A., & Mann, G. E. (2021). Short-chain fatty acids as modulators of redox signaling in health and disease. Redox Biology, 47, 102165. 10.1016/j.redox.2021.102165

Gotoh, M., Sano-Maeda, K., Murofushi, H., & Murakami-Murofushi, K. (2012). Protection of Neuroblastoma Neuro2A Cells from Hypoxia-Induced Apoptosis by Cyclic Phosphatidic Acid (cPA). PLoS ONE, 7(12), e51093. 10.1371/journal.pone.0051093

Guo, Q., Kawahata, I., Degawa, T., Ikeda-Matsuo, Y., Sun, M., Han, F., & Fukunaga, K. (2021). Fatty Acid-Binding Proteins Aggravate Cerebral Ischemia-Reperfusion Injury in Mice. Biomedicines, 9(5), 529. 10.3390/biomedicines9050529

Hamilton, H. L., Kinscherf, N. A., Balmer, G., Bresque, M., Salamat, S. M., Vargas, M. R., & Pehar, M. (2023). FABP7 drives an inflammatory response in human astrocytes and is upregulated in Alzheimer’s disease. GeroScience. 10.1007/s11357-023-00916-0

Huang, W., Guo, H.-L., Deng, X., Zhu, T.-T., Xiong, J.-F., Xu, Y.-H., & Xu, Y. (2017). Short-Chain Fatty Acids Inhibit Oxidative Stress and Inflammation in Mesangial Cells Induced by High Glucose and Lipopolysaccharide. Experimental and Clinical Endocrinology & Diabetes, 125(02), 98–105. 10.1055/s-0042-121493

Islam, A., Kagawa, Y., Miyazaki, H., Shil, S. K., Umaru, B. A., Yasumoto, Y., Yamamoto, Y., & Owada, Y. (2019). FABP7 Protects Astrocytes Against ROS Toxicity via Lipid Droplet Formation. Molecular Neurobiology, 56(8), 5763–5779. 10.1007/s12035-019-1489-2

Jaworska, J., Ziemka-Nalecz, M., Sypecka, J., & Zalewska, T. (2017). The potential neuroprotective role of a histone deacetylase inhibitor, sodium butyrate, after neonatal hypoxia-ischemia. Journal of Neuroinflammation, 14(1), 34. 10.1186/s12974-017-0807-8

Kato, T., Yoshioka, H., Owada, Y., & Kinouchi, H. (2020). Roles of fatty acid binding protein 7 in ischemic neuronal injury and ischemia-induced neurogenesis after transient forebrain ischemia. Brain Research, 1736, 146795. 10.1016/j.brainres.2020.146795

Kim, H., Kim, S., Chung, A.-Y., Bae, Y.-K., Hibi, M., Lim, C. S., & Park, H.-C. (2008). Notch-regulated perineurium development from zebrafish spinal cord. Neuroscience Letters, 448(3), 240–244. 10.1016/j.neulet.2008.10.072

Kim, S. Y., Chae, C. W., Lee, H. J., Jung, Y. H., Choi, G. E., Kim, J. S., Lim, J. R., Lee, J. E., Cho, J. H., Park, H., Park, C., & Han, H. J. (2020). Sodium butyrate inhibits high cholesterol-induced neuronal amyloidogenesis by modulating NRF2 stabilization-mediated ROS levels: involvement of NOX2 and SOD1. Cell Death & Disease, 11(6), 469. 10.1038/s41419-020-2663-1

Lee, J., d’Aigle, J., Atadja, L., Quaicoe, V., Honarpisheh, P., Ganesh, B. P., Hassan, A., Graf, J., Petrosino, J., Putluri, N., Zhu, L., Durgan, D. J., Bryan, R. M., McCullough, L. D., & Venna, V. R. (2020). Gut Microbiota–Derived Short-Chain Fatty Acids Promote Poststroke Recovery in Aged Mice. Circulation Research, 127(4), 453–465. 10.1161/CIRCRESAHA.119.316448

Lee, Y., Lee, S., Park, J.-W., Hwang, J.-S., Kim, S.-M., Lyoo, I. K., Lee, C.-J., & Han, I.-O. (2018). Hypoxia-Induced Neuroinflammation and Learning–Memory Impairments in Adult Zebrafish Are Suppressed by Glucosamine. Molecular Neurobiology, 55(11), 8738–8753. 10.1007/s12035-018-1017-9

Li, H., Sun, J., Wang, F., Ding, G., Chen, W., Fang, R., Yao, Y., Pang, M., Lu, Z.-Q., & Liu, J. (2016). Sodium butyrate exerts neuroprotective effects by restoring the blood-brain barrier in traumatic brain injury mice. Brain Research, 1642, 70–78. 10.1016/j.brainres.2016.03.031

Liu, L., Huh, J. R., & Shah, K. (2022). Microbiota and the gut-brain-axis: Implications for new therapeutic design in the CNS. EBioMedicine, 77, 103908. 10.1016/j.ebiom.2022.103908

Livak, K. J., & Schmittgen, T. D. (2001). Analysis of Relative Gene Expression Data Using Real-Time Quantitative PCR and the 2−ΔΔCT Method. Methods, 25(4), 402–408. 10.1006/meth.2001.1262

Matsumata, M., Inada, H., & Osumi, N. (2016). Fatty acid binding proteins and the nervous system: Their impact on mental conditions. Neuroscience Research, 102, 47–55. 10.1016/j.neures.2014.08.012

Mirzaei, R., Bouzari, B., Hosseini-Fard, S. R., Mazaheri, M., Ahmadyousefi, Y., Abdi, M., Jalalifar, S., Karimitabar, Z., Teimoori, A., Keyvani, H., Zamani, F., Yousefimashouf, R., & Karampoor, S. (2021). Role of microbiota-derived short-chain fatty acids in nervous system disorders. Biomedicine & Pharmacotherapy, 139, 111661. 10.1016/j.biopha.2021.111661

Needham, H., Torpey, G., Flores, C. C., Davis, C. J., Vanderheyden, W. M., & Gerstner, J. R. (2022). A Dichotomous Role for FABP7 in Sleep and Alzheimer’s Disease Pathogenesis: A Hypothesis. Frontiers in Neuroscience, 16. 10.3389/fnins.2022.798994

Offermanns, S. (2014). Free Fatty Acid (FFA) and Hydroxy Carboxylic Acid (HCA) Receptors. Annual Review of Pharmacology and Toxicology, 54(1), 407–434. 10.1146/annurev-pharmtox-011613-135945

Robles-Vera, I., Toral, M., de la Visitación, N., Aguilera-Sánchez, N., Redondo, J. M., & Duarte, J. (2020). Protective Effects of Short-Chain Fatty Acids on Endothelial Dysfunction Induced by Angiotensin II. Frontiers in Physiology, 11. 10.3389/fphys.2020.00277

Rose, S., Bennuri, S. C., Davis, J. E., Wynne, R., Slattery, J. C., Tippett, M., Delhey, L., Melnyk, S., Kahler, S. G., MacFabe, D. F., & Frye, R. E. (2018). Butyrate enhances mitochondrial function during oxidative stress in cell lines from boys with autism. Translational Psychiatry, 8(1), 42. 10.1038/s41398-017-0089-z

Rui, Q., Ni, H., Lin, X., Zhu, X., Li, D., Liu, H., & Chen, G. (2019). Astrocyte-derived fatty acid-binding protein 7 protects blood-brain barrier integrity through a caveolin-1/MMP signaling pathway following traumatic brain injury. Experimental Neurology, 322, 113044. 10.1016/j.expneurol.2019.113044

Sasso, J. M., Ammar, R. M., Tenchov, R., Lemmel, S., Kelber, O., Grieswelle, M., & Zhou, Q. A. (2023). Gut Microbiome–Brain Alliance: A Landscape View into Mental and Gastrointestinal Health and Disorders. ACS Chemical Neuroscience, 14(10), 1717–1763. 10.1021/acschemneuro.3c00127

Storch, J., & Corsico, B. (2023). The Multifunctional Family of Mammalian Fatty Acid– Binding Proteins. Annual Review of Nutrition, 43(1), 25–54. 10.1146/annurev-nutr-062220-112240

Sun, Y., Zhang, H., Zhang, X., Wang, W., Chen, Y., Cai, Z., Wang, Q., Wang, J., & Shi, Y. (2023). Promotion of astrocyte-neuron glutamate-glutamine shuttle by SCFA contributes to the alleviation of Alzheimer’s disease. Redox Biology, 62, 102690. 10.1016/j.redox.2023.102690

Thomas, S. P., & Denu, J. M. (2021). Short-chain fatty acids activate acetyltransferase p300. ELife, 10. 10.7554/eLife.72171

van de Wouw, M., Boehme, M., Lyte, J. M., Wiley, N., Strain, C., O’Sullivan, O., Clarke, G., Stanton, C., Dinan, T. G., & Cryan, J. F. (2018). Short-chain fatty acids: microbial metabolites that alleviate stress-induced brain–gut axis alterations. The Journal of Physiology, 596(20), 4923–4944. 10.1113/JP276431

Weinberg, J. M., Venkatachalam, M. A., Roeser, N. F., & Nissim, I. (2000). Mitochondrial dysfunction during hypoxia/reoxygenation and its correction by anaerobic metabolism of citric acid cycle intermediates. Proceedings of the National Academy of Sciences, 97(6), 2826–2831. 10.1073/pnas.97.6.2826

Zhang, Q., Schepis, A., Huang, H., Yang, J., Ma, W., Torra, J., Zhang, S.-Q., Yang, L., Wu, H., Nonell, S., Dong, Z., Kornberg, T. B., Coughlin, S. R., & Shu, X. (2019). Designing a Green Fluorogenic Protease Reporter by Flipping a Beta Strand of GFP for Imaging Apoptosis in Animals. Journal of the American Chemical Society, 141(11), 4526–4530. 10.1021/jacs.8b13042

Zhang, S., Chen, X., Yang, Y., Zhou, X., Liu, J., & Ding, F. (2011). Neuroprotection against cobalt chloride-induced cell apoptosis of primary cultured cortical neurons by salidroside. Molecular and Cellular Biochemistry, 354(1–2), 161–170. 10.1007/s11010-011-0815-4

