## Supplemental Figure 1 (S1) for "The Neuroprotective Effect of Short-chain Fatty Acids Against Hypoxia-reperfusion Injury"

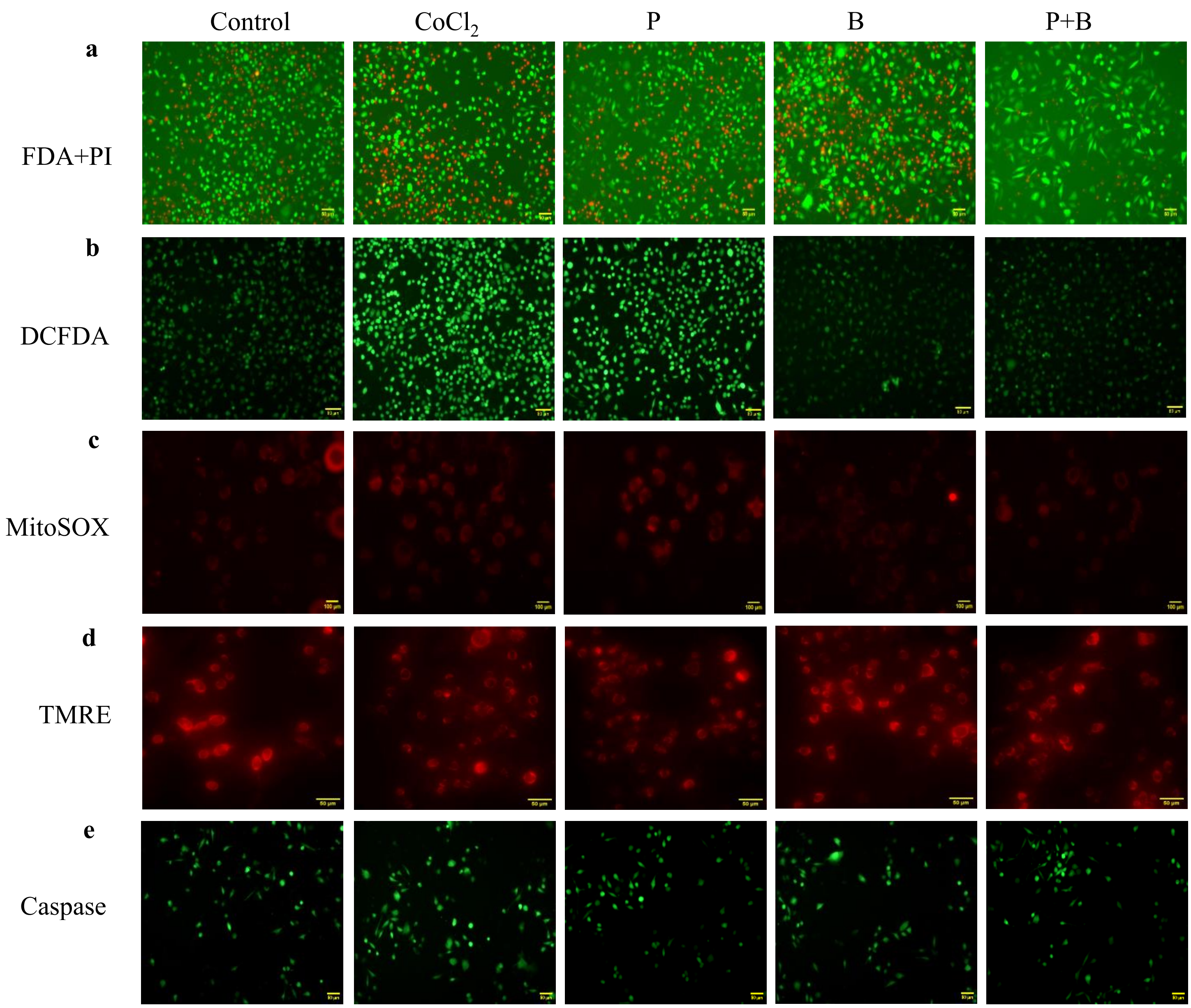


**Fig. S1**. Representative fluorescence images of cells demonstrate the effect of SCFAs on various parameters associated with hypoxia-reperfusion injury. SCFAs mitigate oxidative stress, stabilize mitochondrial membrane potential, inhibit caspase-3 activation, and prevent cell death induced by hypoxia-reperfusion. **a**. FDA-PI double-stained Neuro-2a cells: Live cells are stained with FDA, showing green fluorescence, while dead cells take up PI, represented by red fluorescence. **b**. Intracellular ROS levels, visualized with DCFDA: Increased green fluorescence intensity indicates higher ROS levels observed in CoCl2-treated cells, which decrease upon SCFAs treatment. **c**. Mitochondrial superoxide levels, visualized with MitoSOX Red: Superoxide levels are indicated by increased red fluorescence caused by CoCl2 treatment. **d**. TMRE-stained mitochondria: Loss of $\Delta\Psi m$ is indicated by decreased red fluorescence in CoCl2-treated cells, which is not observed in B and P+B treated cells.
e. Caspase-3 activation, monitored with FlipGFP: The greener the fluorescence intensity, the greater the caspase-3 activation.
